# Zinc finger antiviral protein (ZAP) activity in mammalian and avian hosts in CpG and UpA-mediated restriction of RNA viruses and investigation of ZAP-mediated shaping of host transcriptome compositions

**DOI:** 10.1101/2021.11.04.467232

**Authors:** Valerie Odon, Steven R Fiddaman, Adrian L. Smith, Peter Simmonds

## Abstract

The ability of zinc finger antiviral protein (ZAP) to recognise and respond to RNA virus sequences with elevated frequencies of CpG dinucleotides has been proposed as a functional part of the vertebrate innate immune antiviral response. It has been further proposed that ZAP activity shapes compositions of cytoplasmic mRNA sequences to avoid self-recognition, particularly mRNAs for interferons (IFNs) and IFN-stimulated genes highly expressed when ZAP is upregulated during the antiviral state. We investigated the ZAP functional activity in different species of mammals and birds, and potential downstream effects of differences in CpG and UpA dinucleotide representations in host transcriptomes and in RNA viruses that infect them. Cell lines from different bird orders showed variability in restriction of influenza A virus and echovirus 7 replicons with elevated CpG frequencies and none restricted UpA-high mutants, in marked contrast to mammalian cell lines. Given this variability, we compared CpG and UpA representation in coding regions of ISGs and IFNs with the total cellular transcriptome to determine whether differences in ZAP activity shaped dinucleotide compositions of highly expressed genes during the antiviral state. While type 1 IFN genes typically showed often profound suppression of CpG and UpA frequencies, there was no over-suppression of CpGs or UpAs in ISGs in any species, irrespective of underlying ZAP activity. Similarly, mammalian and avian RNA virus genome sequences were compositionally equivalent as were IAV serotypes recovered from ducks, chickens and humans. Overall, we found no evidence for host variability in ZAP function impacting compositions of antiviral genes.

## INTRODUCTION

Cellular innate immune responses are typically targeted towards pathogen-associated molecular patterns (PAMPs) that enable cellular recognition pathways to differentiate infecting agents from cellular components. Typical PAMPs displayed by viruses that are recognised in vertebrate cells include cytoplasmic double stranded RNA or DNA sequences, non-methylated CpG dinucleotides in viral DNA and abnormally terminated (uncapped and non-poly adenylated) RNA sequences (1, 2). Alternatively, viruses may be recognised through their possession of conserved virus capsid structures, such as the nucleocapsids of retroviruses targeted by TRIM proteins, or complexes formed during virus budding by tetherin (3–6). Recently, selective binding of RNA sequences enriched for CpG dinucleotides by zinc finger antiviral protein (ZAP) was described (7) and this form of compositionally abnormal RNA represents a potential PAMP. ZAP binding triggers antiviral mechanisms that potently restrict replication of RNA viruses and retroviruses with elevated frequencies of CpG in their genomes (8–13). ZAP-dependent restriction of viruses with elevated frequencies of UpA has been similarly characterised (11, 13) and may be mediated through overlapping pathways. While the structural basis of ZAP-mediated restriction of CpG-high sequences has been partially characterised, there appear to be multiple mechanisms downstream that restrict virus replication, including a dependence on TRIM25 (7, 13–16), activation of the nuclease KHNYN (8), a potentially non-canonical activation of oligoadenylate synthetase and its downstream RNAseL RNA degradation pathway (11), or through effects on translation initiation from the bound RNA template (17–19).

The ability of ZAP to identify and restrict replication of viruses with high CpG and UpA frequencies is contingent on the pervasive suppression of both dinucleotides in vertebrate cellular mRNA sequences that remain below the ZAP “radar”. Frequencies of CpG in vertebrate mRNAs range from 20%-80% of expected frequencies based on the frequencies of their component bases (20); suppression is particularly pronounced in mammals and birds and in sequences with a low G+C content (21). Methylation of vertebrate genomic DNA and mutational loss of methylated CpG (22, 23) (and a consequent secondary loss of UpA; (24)) may account for dinucleotide suppression within encoded cellular genes. However, there are consistent differences in degrees of suppression between coding and non-coding RNAs and relationships between CpG and Up frequencies with G+C composition that imply the existence of further selection mechanisms against both dinucleotides in sequences expressed as RNA in the cytoplasm (25).

While the phenomenon of ZAP-mediated restriction of viruses mutated with high CpG or UpA inserts is well characterised, the functional role of this pathway in restricting virus replication in nature is less well understood. Most vertebrate RNA viruses display similar degrees of CpG and UpA suppression to those of the host cells they infect (25–29) rendering the pathway seemingly pointless in the majority of cases. The main exceptions are double-stranded RNA viruses, such as members of the *Reoviridae*, that show little suppression of CpG and may therefore be able to avoid ZAP-mediated restriction through currently uncharacterised evasion or shielding mechanism.

The other prominent exceptions are members of the vector-borne *Alphavirus* genus of the family *Togaviridae*, such as Sindbis virus, which shows much higher CpG frequencies than other RNA viruses of similar G+C content. This has been shown to enable ZAP-mediated restriction of wild type virus in mammalian cells (19, 30, 31). We have proposed incomplete suppression of CpG may represent an “adaptive compromise” for these and other groups of vector-borne viruses, such as flaviviruses (32). While the replication of high CpG mutants of Zika virus was restricted in mammalian cell culture, they replicated to higher titres in insect cell culture and showed greater systemic spread and excretion in saliva in experimentally infected mosquitoes (32). Flaviviruses that replicate in insects only (insect-specific flaviviruses; ISFs) show no equivalent CpG suppression (33), mirroring the absence of methylation and CpG suppression in insect and many other arthropod groups. This supports the idea that dual host-specificity places conflicting evolutionary pressures on genome composition, a problem that may be resolved differently in different virus groups. Alphaviruses appear to have adopt a CpG high genome to facilitate replication in mosquitoes, while vector-borne flaviviruses may be better optimised for replication in vertebrate hosts. The remarkable observation that the apparent host range restriction of ISFs to insect cells may be overcome in mammalian cells by knocking out ZAP expression (34) underlines the involvement of this pathway in determining host range. A specific role of ZAP in preventing transmission of arthropod viruses that do not suppress CpG to mammals and birds remains a speculative possibility; if so, it would be a protective mechanism that vector-borne viruses have able to breach, albeit at some evolutionary cost.

It has been previously proposed that in addition to controlling RNA virus replication, ZAP and allied antiviral restriction pathways may play roles in host gene regulation (35). For example, the greater than expected suppression of CpG frequencies in human type I interferon (IFN) genes (encoding IFN-α, IFN-β) was proposed as a mechanism to enhance expression of these critical antiviral proteins (35). The IFN-inducibility of the long isoform of ZAP (36, 37) may indeed induce a hostile internal milieu in cells challenged by virus infection where high levels of ZAP expression may suppress global gene expression and bias gene expression towards genes with over-suppressed CpG frequencies, such as those for IFNs and thereby enhancing the cellular anti-viral state. Shaw *et al.* have indeed has recently proposed that interferon stimulated genes (ISGs) may also be considered as part of a compositionally distinct “interferome” with suppressed frequencies of CpG that might enable prolonged gene expression after induction of ZAP (38). Experimentally, the hypothesis is difficult to substantiate since CpG is almost universally suppressed in vertebrate transcriptomes and ZAP or ZAP-like proteins have been detected in the wide range of vertebrates investigated to date (39). Recently, however, it was reported that functional suppression of viruses with high CpG frequencies was impaired in cells expressing a range of avian-derived ZAP genes compared to those expressing human ZAP (40). The possibility that avian cells may not suppress replication of high CpG RNA sequences as effectively as mammalian cells was supported by Greenbaum *et al.* (41), who observed that CpG representation in the “Spanish flu” pandemic strain of influenza A virus (IAV) genomes reduced systematically in the decades after 1918 following its zoonotic transfer from birds to humans. This change may have been in response to a more restrictive environment in mammalian cells mediated at least in part by a functional ZAP.

If avian ZAP shows a reduced ability to recognise and suppress expression of mRNAs or inhibit replication of RNA viruses with high CpG RNA sequences, then these functional differences should drive differences in genome compositions between viruses adapted for replication in mammals and birds. Such differences may extend to host-encoded ISG and IFN gene compositions. In the current study, we have detected differences in the restriction of replication of high CpG mutants of influenza A virus and of enterovirus 7 replicons in *Gallus gallus* (chicken)-derived cell lines compared to those of *Anas platyrhynchos* (mallard duck) and other anseriforms, as well as a range of mammalian cell lines. Despite these differences, we detected no systematic differences in CpG or UpA representation in mRNAs, ISGs or in the genomes of viruses infecting chickens, ducks or humans. We therefore obtained no support for the hypothesis that the composition of the cellular transcriptome is conditioned and regulated by ZAP through systematic differences in CpG composition. The similarity in CpG and UpA compositions of IAV strains infecting birds and humans similarly provides little evidence that CpG content is associated with mammalian and avian host range adaptation of RNA viruses.

## MATERIALS AND METHODS

### Cell culture provenance and maintenance

As standard, all cells were cultured at 37°C, in 5% CO_2_ and harvested with TryPLE dissociation reagent (Gibco) for 5 min, centrifugation at 1000 rpm for 5 min followed by resuspension in fresh media and re-seeding into fresh flasks.

Zebra finch cell Gray 266 (G266) were derived from a naturally occurring tumour of a male zebra finch forehead (Itoh and Arnold, 2011) and were kindly donated by Dr Butter Falk. The cells were maintained in DMEM media supplemented with 10% heat-inactivated foetal calf serum, 2% heat-inactivated chicken serum, 36.7 mmol/L of glucose and penicillin and streptomycin (Pen/Strep). The cells were replated at a dilution of one-third. Quail cell line QT6, derived from a Japanese quail fibrosarcoma, were purchased from ECACC. The cells were maintained in Ham’s 10 buffer supplemented with 2mM Glutamine, 10% tryptose broth, 10% foetal calf serum and Pen/Strep. The cells were split at 70 to 80 % confluency and re-plated at one in two dilutions.

Human A549 (purchased from ATCC), A549 ZAP−/−/− (CRISPR engineered from WT A549 cells), chicken DF-1, Hek293T, MDBK, zzR-127, BFA, FBT, AK-D and the primary cell lines for pigeon, partridge and - pheasant were maintained in DMEM-Ham(F12) media supplemented with 10% FCS, 2mM glutamine, 1% non-essential amino acids and 1% P/S. For YO cell line the media included also 1% sodium pyruvate. They were diluted one in ten when re-plating.

Duck embryo cell line CCL-141 (ATCC) was maintained in EMEM medium supplemented with 10% FCS, 1% P/S and subcultured at 70-80% confluency. They were diluted one in five when re-plating. FBT and pig lungs primary cell lines were cultured in similar media with addition of nystatin.

Primary cell lines from pigeon, partridge, pig lung and pheasant were supplied by Animal and Plant Health Agency England. zzR-127, YO and FBT were sourced from the Pirbright institute cell bank. AK-D feline cell line was a kind gift from Pr Ian Goodfellow. BFA was a kind gift from Pr Rick Randall. DF-1, Hek293T and MDBK cell lines were from personal cell collection of the author’s cell bank.

### E7 replicon assay

The echovirus 7 (E7) replicon was based on the Wallace strain and modified by insertion of compositionally modified sequences in the 5’UTR as previously described (42).

### RNA transcription and Transfection

T7 transcription reactions (MEGAscript T7 transcription kit-Promega) were assembled according to manufacturer’s recommendation with 1 μg of E7 plasmid DNA linearised with *Not*I or TK-Ren plasmid linearised with *Xba*I. Following a 2 h incubation, and DNAse treatment, the transcripts were purified using RNA clean and concentrator kit (Zymo Research) and the concentration measured using the Qubit fluorometric quantitation system (ThermoFisher Scientific). 50 ng of E7 RNA was transfected to each well together with 5 ng of renilla RNA. The transfection mix was prepared using Lipofectamine MessengerMax (Invitrogen) and OptiMem (Gibco). At 6 h post transfection, the media was discarded and the wells were washed with PBS before adding 50 μl of Passive lysis buffer (Promega). Cells were frozen at −20°C to facilitate complete lysis.

### Determination of luciferase activities

Luciferase activities were determined using a GloMax Multi detection system (Promega) reader. Each well was injected with Dual Luciferase reagents (Promega). This method consists of first injecting 50 μl of firefly luciferase reagent to an equal volume of cell lysates. Following a 30 min period for the complete decay of the luminescence, 50μl of a stop/renilla reagent was injected to measure the renilla luminescence in the same well. Firefly luminescence was normalised using the renilla values. The experiments were carried out in 3 biological replicates with 4 technical replicates.

### Reconstitution of chicken cell line with human ZAP

A549 ZAP^−/−/−^ and DF-1 cells were cultured in 10 cm dish and transfected using LT1 reagent (Mirus lab) with 5 μg of huZAP-GFP DNA for 24 h. The cells were transfected with 5 μg of E7 replicons WT, CpG-H and 500 ng of TKRen RNA using Lipofectamine MessengerMax. Following 6 h incubation, the cells were trypsinised, resuspended in 1% FCS-PBS buffer and sorted according to green fluorescence using a BDFACS ARIAIII machine sorter to isolate GFP expressing cells. 30,000 cells were collected in passive lysis buffer in white 96 well plate and the luciferase activities were determined as previously described.

### Cloning of IAV mutants

The WSN system was elaborated by Hoffman and colleagues (43) and consists of 8 pHW2000 expression plasmids each encoding one of the viral segments from the H6N1 strain of influenza A virus (IAV A) and driven by CMV promoter. The system allows manipulation of individual segments of the virus.

The HA segment (segment 4) was selected as it is the most variable segment and contains only one long open reading frame of 1775bp. The first 171 and last 107 bases were not muatenised. The mutations introduced into the rest of the HA segment maintained the WT amino acid sequence while increasing the CpG or UpA content were designed using the SSE software package (44), sequences listed in Table S1; Suppl. Data). The observed to expected (O/E) ratio of CpG in WT of 0.38 was increased to 1.9 in CpG high (CpG-H) segment. The mutated segments were ordered from GeneArt (Thermo Fisher). The segments were amplified by PCR using KOD polymerase creating blunt-end products (Merck) using primers listed in Table S2 (Suppl. Data). The primers were designed to create overlapping ends with the wild type sequence of the pHW244-HA vector. The PCR products were *Dpn*I treated and gel purified before proceeding with Gibson assembly.

The Gibson assembly was performed with the NEBuilder HiFi DNA assembly (NEB) using 50 ng of vector PCR and 2-fold excess of insert PCR (CpG-H) following the manufacturer’s instructions. The reaction mixes were incubated at 50 °C for 30 min. 2 μl of assembled plasmids were transformed in competent TOP10 *E. coli* cells and selected on LB/ampicillin agar plates. The integrity of the mutant plasmids was verified by Sanger sequencing using primer pairs ‘S4 CGH insert F + R’ (Table S2; Suppl. Data).

### Rescue of IAV

The transfection reagent was prepared as follow: in one tube, 0.5 μg of DNA for each of the eight plasmids constituting the WSN system were mixed with 25 μl OptiMEM. For the negative control, one of the plasmid (the HA coding sequence) was omitted. In the second tube, 6 μl of Lipofectamine 2000 were mixed with 125 μl of OptiMEM. Both tubes were then combined and incubated for 20 min at room temperature.

A confluent T75 flask of HEK 293T cells was trypsinised and harvested by centrifugation and resuspended in 5 ml of DMEM 10% FCS, 1% pen/strep. Five hundred microlitres of the cell suspension were then seeded in a 6 well-plate and mixed with the DNA transfection mix. After 6 h the media was changed to DMEM 0.5% FCS, 1% pen/strep medium. The supernatant was collected following a 48 h incubation period. The viral supernatant was centrifuged at 3000 rpm for 3 min and aliquots were stored at −80 °C.

### Plaque assay

To resolve the viral titre of recued viruses 300,000 MDBK cells were seeded per well of 12-well plates. The following day, a 10-fold serial dilution of the virus stock was prepared with extensive mixing by vortexing between dilutions. The media was aspirated from each well. The cells were washed with PBS, and each well was inoculated with 150 μl of diluted virus. The plates were incubated at room temperature for 1 h with regular rocking for viral absorption. The inoculum was discarded, and wells were washed with PBS. A 1% low melting point agarose (GeneFlow) / 0.5% FCS-DMEM overlay was added to each well, following a 15 min setting time at room temperature, the plates were returned to an incubator inverted for 3 days. Plaques stained with crystal violet were visible as clear spots and counted.

### Replication phenotypes with IAV mutants

DF-1, CCL-141 and QT6 were seeded in 6 well plates at 500,000 cells and infected with IAV WT or mutant CpG-H or UpA-H virus at MOI 0.001 for 1h. The inoculum was discarded and the cells were washed with PBS. The cells were cultured in 0.5% FCS-DMEM culture media. At 1, 12, 24, 36, 48, 60 and 72 h post-infection (p.i.) an aliquot was collected and the virus titre determined by TCID_50_ in A549 ZAP^−/−/−^ cells. This was performed with twoindependent duplicate for each virus and cell line.

### Quantitative PCR

Cells were stimulated using poly (I:C). DF-1 cells were stimulated with 10 μg/ml of poly (I:C) added to the media and QT6, pigeon, G266 and CCL-141 cells lines were transfected with 1 μg of poly (I:C) using Lipofectamine 2000. At 0, 2, 4, 6 and 8 h post-stimulation, cells were harvested and RNA was purified using a total RNA purification kit (RNeasy Qiagen). 80 ng of total RNA was added as template for real-time PCR using QuantiFast SYBR green Master Mix (Qiagen) in the StepOne plus instrument (Applied Biosystems). The programme for cDNA synthesis and RNA quantitation was as follow for the Reverse transcription step (50°C for 10 min- 95°C for 5 min) and for quantitation 40 cycles (95°C for 10 sec, 60°C for 30 sec).

Expression of each gene was normalized to an internal control (HPRT1) transcript, and these values were then normalized to the non-stimulated control cells to yield a fold induction value. Primers used for the detection of HPRT1, IFNβ, and ZAP are listed in Table S3 (Suppl. Data).

### Detection of positive selection in ZAP genes

Avian ZAP (encoded by the *ZC3HAV1* gene) sequences were obtained from Ensembl (Table S4; Suppl. Data). Sequences were aligned using Muscle v. 3.8.425 (45) and trimmed to remove the highly diverse region between the fourth and fifth zinc-finger motif. To test for positive selection across sites in the alignment, maximum likelihood analysis of ratios of nonsynonymous to synonymous nucleotide substitutions (dN/dS; ω) was performed with the codeml package of programs in PAML v. 4.9 (46, 47). Various site models were fitted to the multiple alignments: M1a (neutral model; two site classes: 0 < ω_0_ < 1 and ω_1_ = 1); M2a (positive selection; three site classes: 0 < ω_0_ < 1, ω_1_ = 1, and ω_2_ > 1); M7 (neutral model; values of ω fit to a beta distribution where ω > 1 disallowed); M8 (positive selection; similar to M7 but with an additional codon class of ω > 1); and M8a (neutral model; similar to M8 but with a fixed codon class at ω = 1). Likelihood ratio tests were performed on pairs of models to assess whether models allowing positively selected codons gave a significantly better fit to the data than neutral models (model comparisons were M1a vs. M2a, M7 vs. M8, and M8a vs. M8). In situations where the null hypothesis of neutral codon evolution could be rejected (P < 0.05), the posterior probability of codons under selection in M2a and M8 was inferred using the BEB algorithm (48). In addition, to test for positive selection in different branches of the phylogeny, the ‘free-ratios’ model (which fits one ω for every branch in the tree) was implemented in PAML (49).

To calculate the nucleotide diversity in the avian ZAP sequences used for PAML analysis, the alignment was analysed using the PopGenome package in R (50). A sliding window of 100 bp and an increment of 1 bp was used to calculate nucleotide diversity across the alignment, which is expressed as the average pairwise number of variant sites per 100 bp window.

### Analysis of dinucleotide frequencies in host genomes

Non-redundant mRNA sequences of human, chicken and duck were downloaded from the http://www.ncbi.nlm.nih.gov/gene database, with coding region sequences shorter than 450 bases excluded. Mono- and dinucleotide frequencies and ratios of observed dinucleotide frequencies to those expected from mononucleotide composition (G+C content in the case of DNA sequences) were calculated using the program Composition Scan in the SSE package (44). All statistical analyses were performed using SPSS version 26.

## RESULTS

### Functional investigation of CpG- and UpA-mediated restriction of RNA virus replication in avian and mammalian cell lines

The E7 replicon was modified through the insertion of WT, CpG-H, UpA-H, 2xUpA-H and a scramble control (CDLR) sequences into the 3’UTR. Addition of WT sequences in this non-translated region of the replicon had little effect on replication while insertion of sequences with elevated CpG or UpA frequencies lead to a ZAP-dependent attenuation of replication as previously described (11). The effects of compositional modification on replication in the chicken derived DF-1 cell lines was investigated by transfection of 50 ng of *in vitro* transcribed RNA of WT and mutants of the E7 replicon and monitoring replication at 6 hours post-infection as previously described (42). The experiment was performed in parallel with control human cells A549 and a CRISPR derived ZAP knock-out (k/o) A549 cell line, B6 used in previous ZAP functional assays (11).

The CpG-H replicon showed a 10-fold reduction in replication compared to WT in the control human A549 cell line as previously observed (42) and this attenuation was reversed in the ZAP k/o cell line (Fig. 1). Attenuation of the UpA-H and 2xUpH-A replicon mutants and reversion in the k/o cell line was similarly observed. In contrast to A549 WT cells, no attenuation of the CpG-H or either UpA-H replicon was evident in the chicken DF-1 cell line. To determine whether this lack of restriction was the result of impaired or absent expression of chicken ZAP, human ZAP fused with GFP was transfected in DF-1 cells followed by FACS-based sorting to isolate expressing cells based on fluorescence. The cells were then transfected with WT and CpG H replicons. The reconstitution of human ZAP in human k/o A549 cell line showed a small but statistically significant attenuating effect on the CpG-H replicon. However, no attenuation of replication of the CpG-H mutant was observed in the chicken DF-1 cell line irrespective of reconstitution with huZAP (Fig. 1B).

**FIGURE 1.**
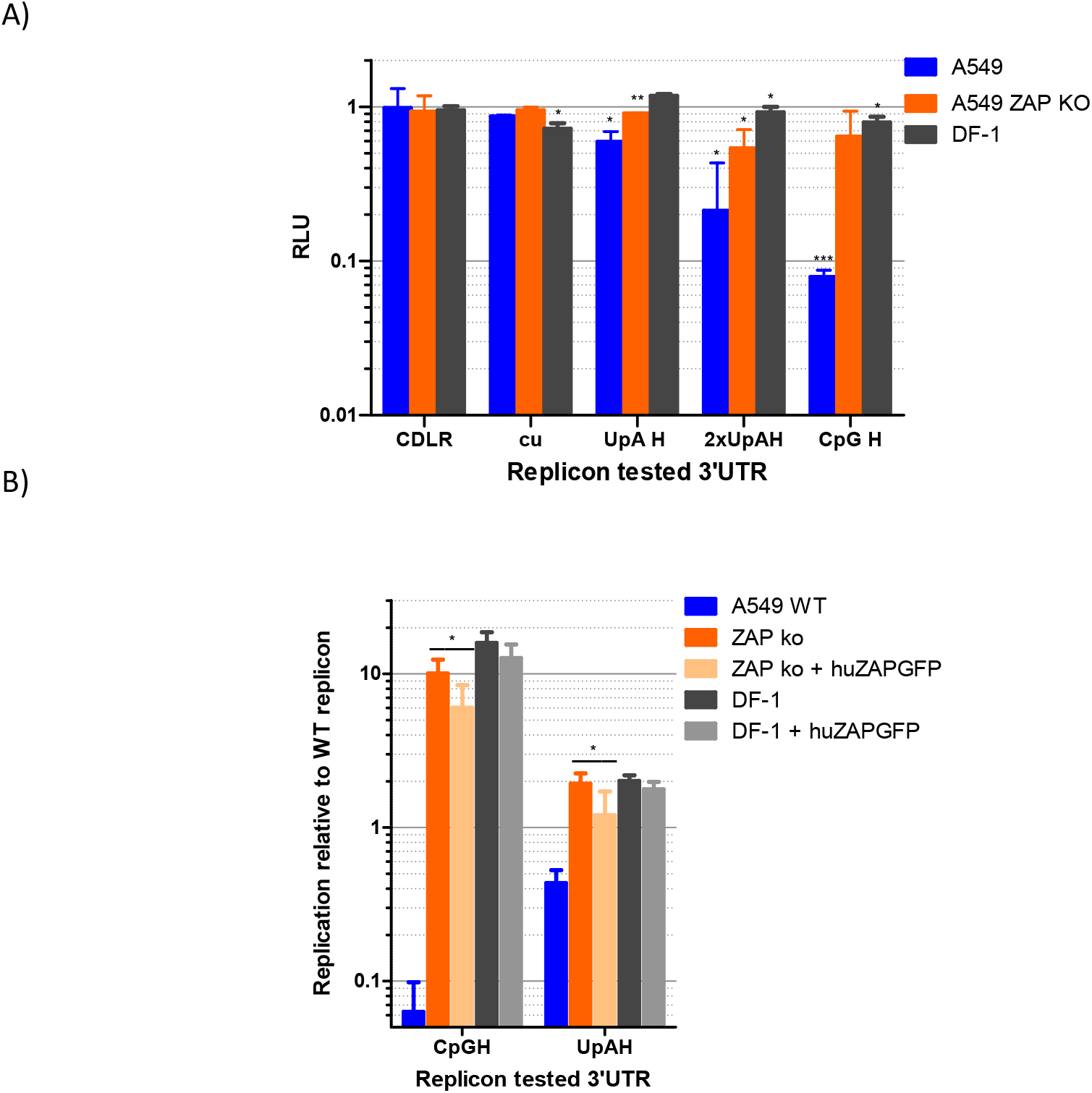
Replication of E7 replicons in control A549 and chicken DF-1 cell line. Cells were transfected with 50ng/well of in vitro transcribed RNA of E7 replicon with a 3’UTR variant WT, permuted (CDLR), CpG low (cu) and elevated of CpG (CpG-H) and UpA (UpA-H) and assayed for luciferase expression at 6h post transfection. (A) Relative replication in control human cell lines A549, A549 ZAP k/o and DF-1 cells. (B) effect of overexpression of human ZAP in A549 ZAP ko and chicken DF-1 cell lines compared to WT. Cells were transfected with a human ZAP fused to GFP to enable FACS sorting of GFP expressing cells and transfected with E7 WT and mutant versions. Significance of differences from WT replication were calculated by two tailed paired t test; asterisks show significance values as follows: ***: *p* < 0.001; **: *p* < 0.01 and *: *p* < 0.05

To investigate whether the DF-1 cells typified avian antiviral responses to E7 and the extent to which cell lines from different mammalian species might vary, we transfected E7 replicons into a range of avian and mammalian cell lines and determined the extent of attenuation of CpG-H and UpA mutants by reference to replication levels in the WT E7 replicon (Fig. 2). As controls, mutants with 3’UTR sequences scrambled by CDLR or with minimised frequencies of CpG and UpA were constructed and tested in parallel. As previously described, replication of both control replicons was comparable to WT replicons in A549 cells and in A549 ZAP k/o cells; replication in each mammalian cells lines was comparable in each of the mammalian cells lines with the exception of the replication of the CDLR mutants in AK-D cells although the cu mutant replicated to similar levels to those seen with the WT replicon. In contrast, the CpG-H replicon was attenuated in all mammalian cell lines tested: rat (YO), mouse (BV2), goat (zzR-127), bovine cell lines (FBT and BFA), cat (AK-D) and primary pig lungs showed reductions in replication of 6-10 fold compared to the WT replicon (Fig. 2A). There was a comparable reduction in the replication of the UpA-high replicon mutants, with the double UpA mutant showing levels of attenuation comparable to that of the CpG high mutant. Attenuation of both UpA-H mutants was reversed in the ZAP k/o cell line, indicative of a direct or indirect role of ZAP in their attenuation.

**FIGURE 2.**
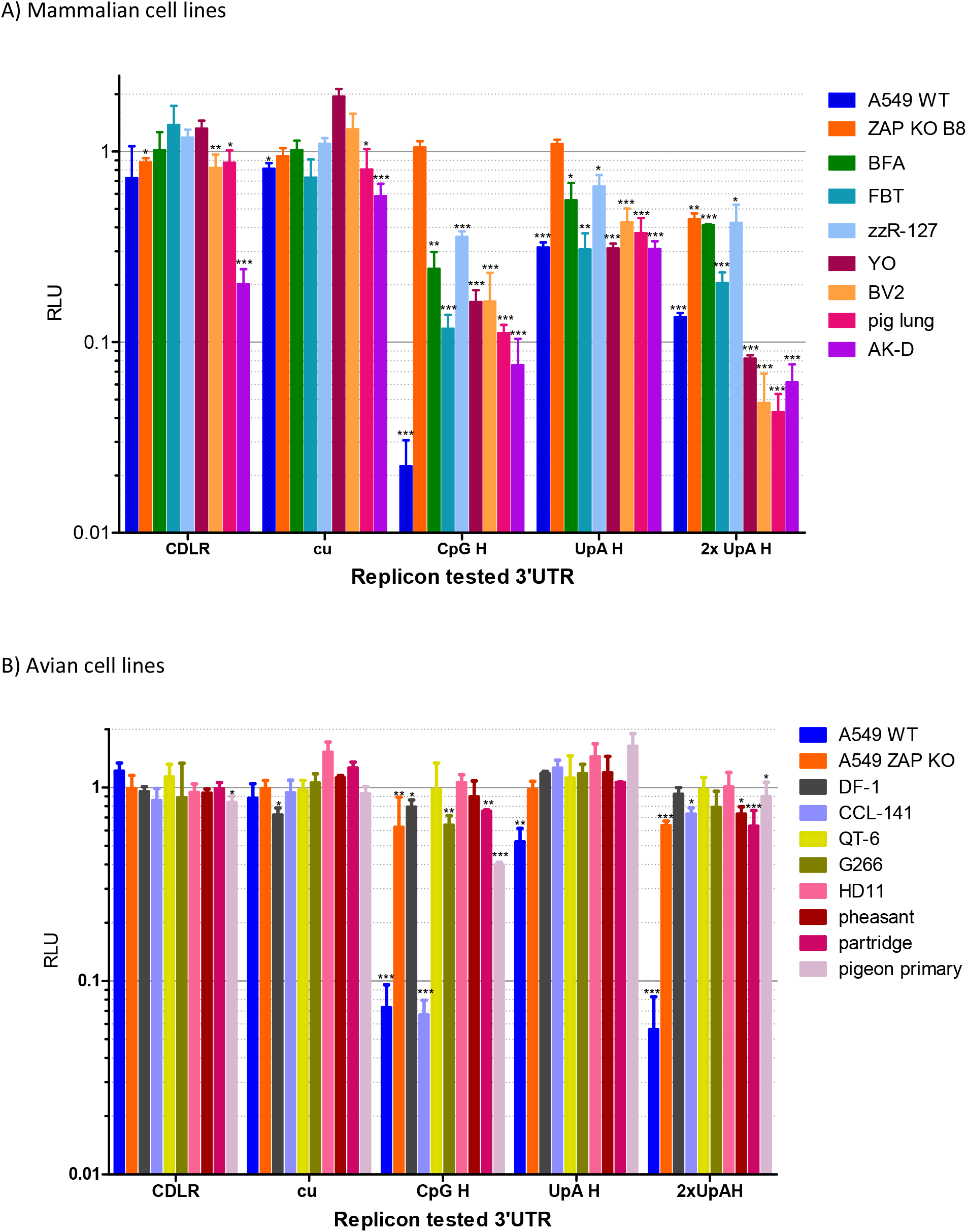
Replication of E7 replicons in a range of (A) mammalian and (B) avian cell lines. Cells were transfected with 50ng/well of in vitro transcribed RNA of E7 replicons as described in Fig. 1. The bar heights depict mean replication relative to the WT replicon in each cell line; the data were derived from 3 biological replicates; error bars show standard deviations. Significance of differences from WT replication were calculated by two tailed paired t test; asterisks show significance values as follows: ***: *p* < 0.001; **: *p* < 0.002 and *: *p* < 0.033. Abbreviations: mammalian cell lines: BFA-Bovine Fetal Aorta; FBT: Foetal Bovine Turbinate; zzR-127: goat foetal tongue cell line; YO: derived from a hybrid myeloma YB2/3HL; BV2: microglial cells; AK-D: Foetal cat lung. Avian call lines: DF-1: chicken embryo fibroblast; CCL-141:-Duck embryo; QT6: Japanese Quail fibrosarcoma; G266: male zebra finch; HD11: chicken macrophage-like; primary cells from pigeon (Columbaves), partridge and quail (Galliformes).

Parallel assay of a range of avian cell lines derived from members of the Galliformes, Anseriformes, Passeriformes and Columbaves orders revealed markedly contrasting restriction of the CpG-H mutant (Fig. 2B). Representing Galliformes, cell lines included the macrophage-like HD11 cell line from chicken, QT6 from quail and primary cell lines from pheasant and partridge. We also evaluated replication in the duck embryo cell line CCL-141 (Anseriformes) and zebra finch G266 cell line (Passeriformes) and pigeon primary cells (Columbaves). There was no attenuation of CpG-H or UpA-H replicons in any of the galliform-derived cells (DF1 fibroblasts and HD11 macrophages from chicken, QT6 cells from Quail or primary embryo fibroblasts from pheasant or partridge). Similarly, there was no inhibition detected in the zebra finch cell line G266. However, the duck CCL-141 cell line and pigeon primary cells exhibited complete or partial restriction of the CpG-H E7 replicon construct. Of note and in marked contrast to mammalian cells, none of the avian cell lines showed any restriction of either UpA-H mutant. This included the duck CCL-141 cell line in which the CpG-H mutant was severely attenuated (Fig 2B).

To verify the apparent difference in restriction of the E7 replicon mutants between avian species in a virus system, we employed an established reverse genetics system for influenza A virus to create mutants of IAV with compositionally modified segment 4 sequences encoding haemagglutinin (HA). A total of 100 additional CpG or UpA dinucleotides were introduced without modifying the amino acid sequence of the encoded HA protein sequence. Modifications were made only the mid-section of this segment to avoid potential unintended effects of mutagenesis on viral transcriptional and packaging signals at the segments ends (whole segment sequences are listed in Table S1; Suppl. Data). The replication kinetics of WT and mutants of IAV from cells infected at low MOI (0.001 / cell) were comparable in the galliform QT6 and DF-1 cell lines from quail and chicken respectively (Fig. 3A). However, while replication of WT IAV in the duck-derived cell line, CCL-141, was comparable to that of other cell lines, the CpG-H mutant was substantially attenuated, producing approximately one log less infectious virus at each collection point over the 72-hour time course. Consistent with the data obtained with the E7 replicon, no attenuation of the UpA mutant IAV was observed (Fig. 3B).

**FIGURE 3.**
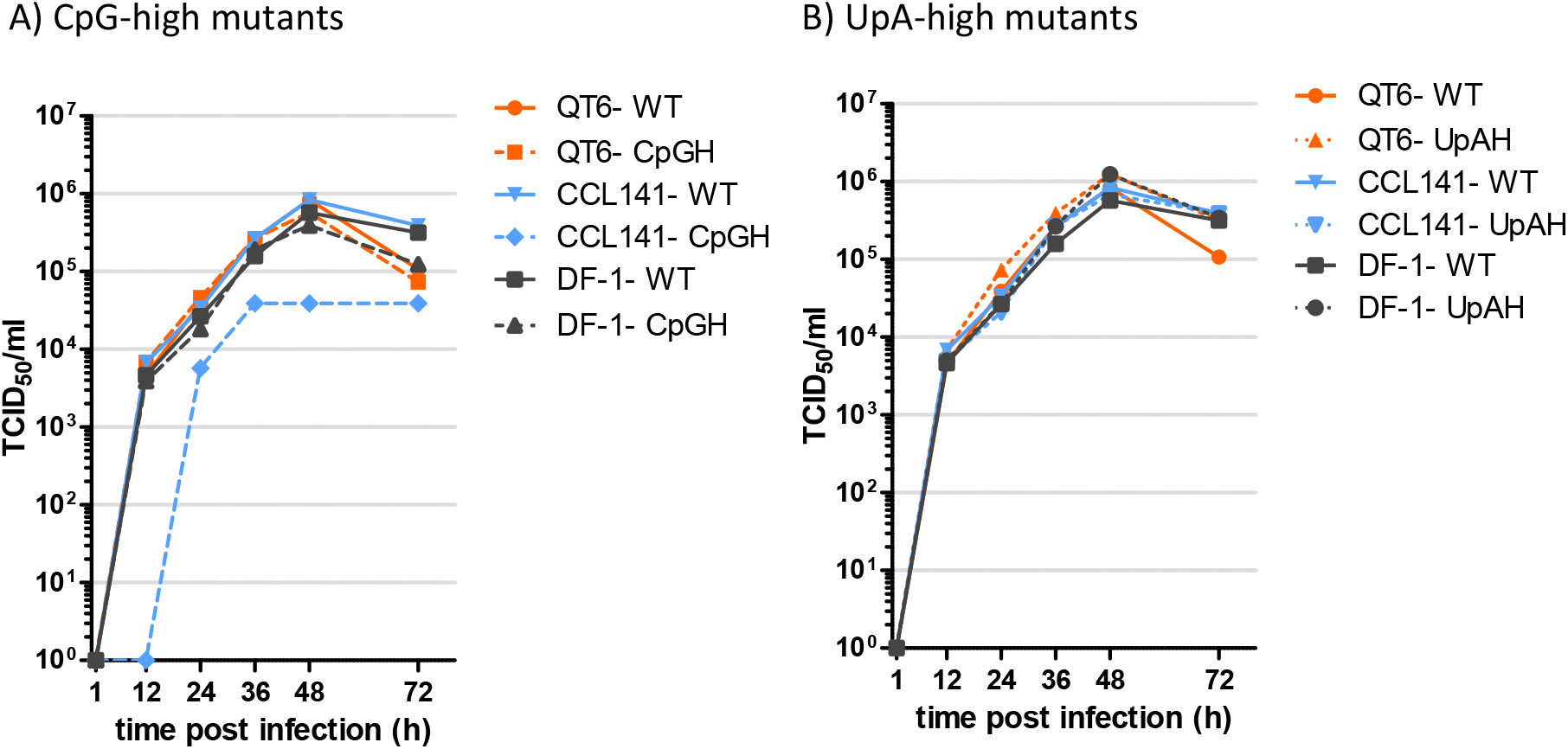
Comparison of the replication kinetics of WT IAV with mutants with elevated (A) CpG and (B) UpA dinucleotide frequencies in segment 4. Replication was assayed in mutants in the avian cell lines QT6, DF-1 and CCL-141. Cells were infected at an MOI of 0.001 and at specified times supernatant were collected and titrated by infectivity using A549 ZAP k/o cells – TCID_50_ values shown on the y-axis. Data points represent the mean of 2 biological replicates.

To investigate whether the lack of CpG-mediated restriction observed in chicken cells was the result of reduced expression or a failure of induction of ZAP on infection, we determined mRNA levels of ZAP and IFN-β in DF-1 cells after poly(I:C) stimulation and compared responses with those in CCL-141 cells (Fig. 4). In both cell lines, poly(I:C) treatment induced expression of IFN-β indicating that cells were responsive and both cell lines upregulated of ZAP mRNA at 8 hours post-stimulation. Based on this comparison, the failure of restriction of CpG-high mutants in DF-1 cells cannot be attributed to a lack of cellular upregulation of ZAP expression. Consistent with this finding, there was a comparable absence of CpG- or UpA-mediated restriction in DF-1 cells pre-treated with poly-I:C (Fig. S1; Suppl. Data).

**FIGURE 4.**
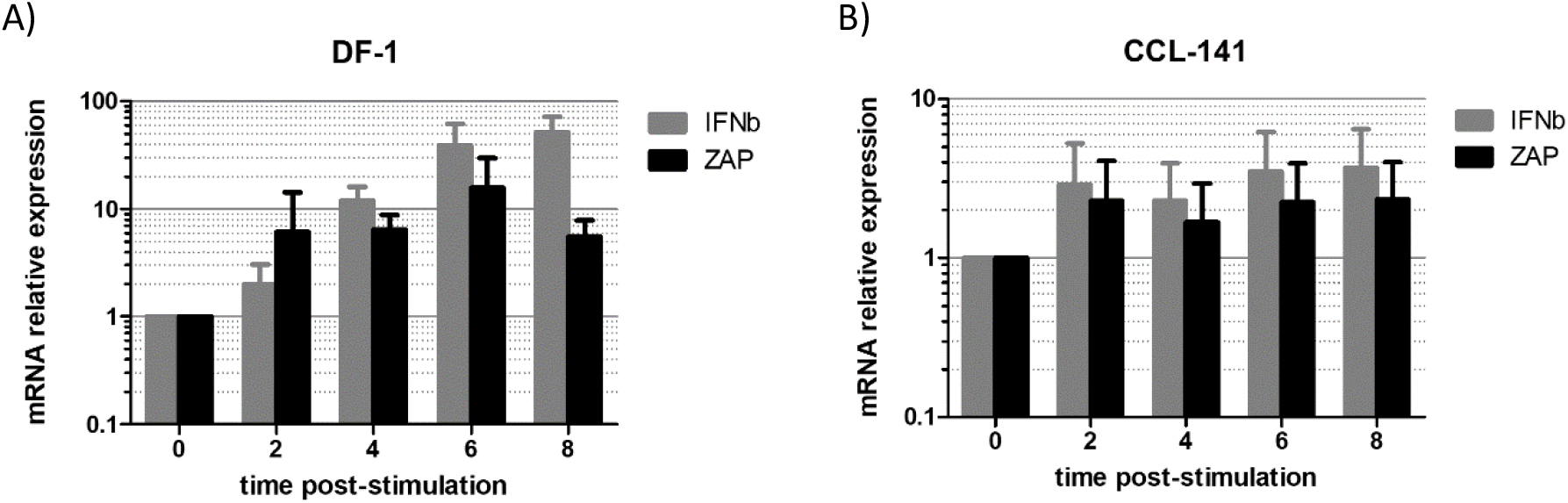
Induction of IFN-β and ZAP after poly(I:C) stimulation. RNA extracted from cell lysates of (A) DF-1 chicken, and (B) CCL-141 duck cell lines at different time points post-stimulation was quantified for ZAP and IFN-β mRNA expression by qRT-PCR. Data shows mean fold-change in expression relative to unstimulated control.

### Adaptive evolution of avian ZAP

Evidence for rapid adaptive changes in ZAP gene organisation and function in the primate lineage has been proposed to indicate the importance of ZAP in host cell defence (51). The variability in ZAP function in avian cell lines observed in the current study may reflect effects of different selection pressures on the evolution of the avian ZAP gene locus. To investigate whether (and along which lineage) avian ZAP has undergone potential virus-driven adaptive evolution, site and branch models were implemented in PAML to detect positive selection. The majority of sites in the alignment were found to be under purifying or neutral selection (87.93%, M2a; 84.17%, M8; Table S5; Suppl. Data), while a smaller proportion (12.07%, M2a; 15.83%, M8) were determined to be under positive selection. Positive selection was highly significant (p ≈ 0) for all site model comparisons, compared to the null hypothesis of purifying and neutral evolution. Using the Bayes Empirical Bayes procedure, codons evolving under positive selection were identified (n = 16 for M2a, n = 27 for M8; Table S5; Suppl. Data). Codons under positive selection tended to cluster around the zinc-finger domain (n = 7, of which 6 were found within the CCCH motif itself) and the PARP domain (n=6). Furthermore, one positively selected site, 527V, was found within the NAD+ binding site in the PARP domain (Fig. 5A).

**FIGURE 5.**
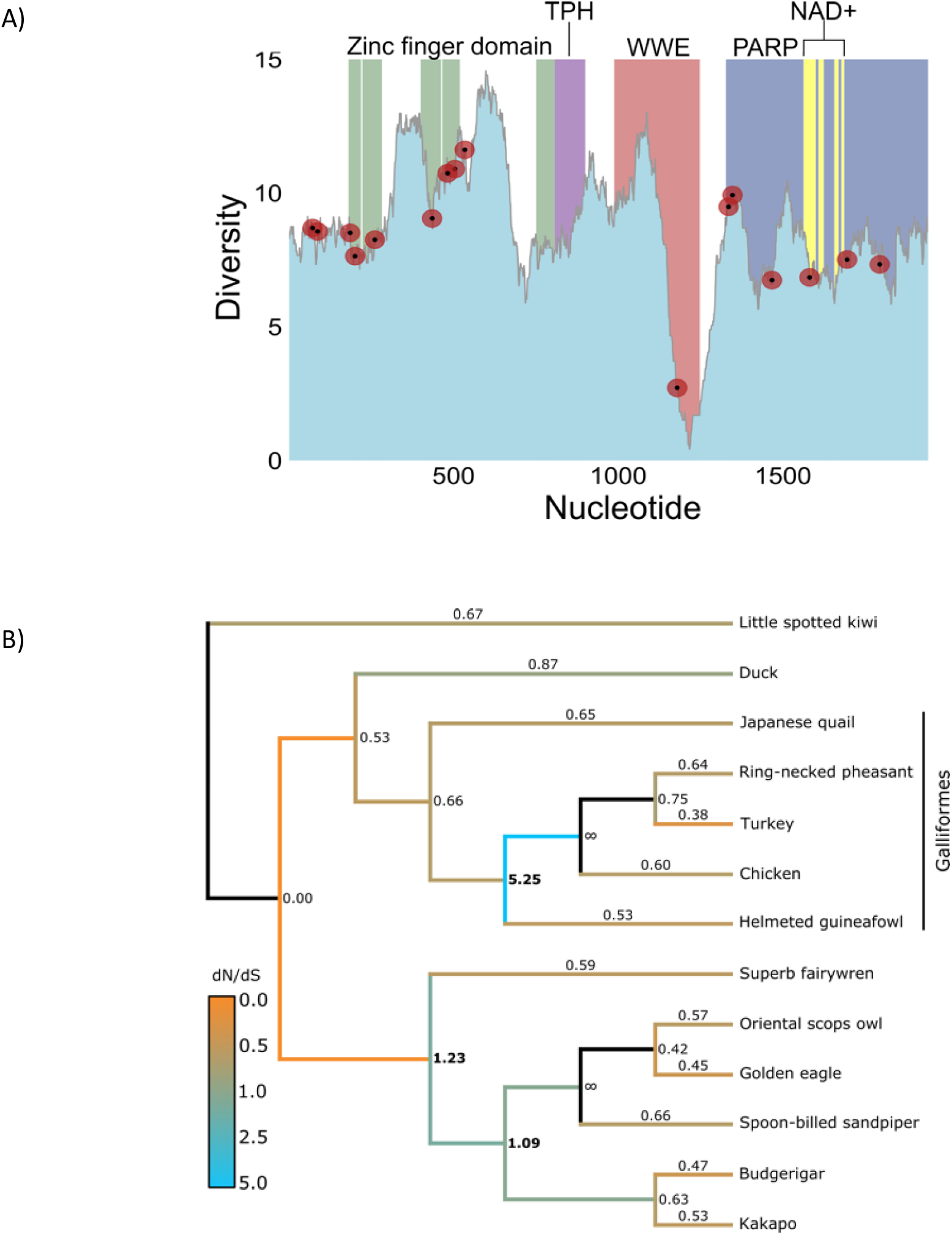
Diversity and positive selection analysis of avian ZAP. (A) Using an alignment of 13 avian ZAP sequences (Table S4A; Suppl. Data) and following trimming to remove a region of no homology between the fourth and fifth zinc-finger motifs, a sliding-window analysis of sequence diversity was conducted using 100bp windows and 1bp increment size. Diversity was calculated as the average pairwise number of variant sites per 100bp window. The four domains of ZAP are shown in coloured blocks, including the NAD+ binding sites in the PARP domain (yellow). Superimposed in red points are positively selected sites as determined by M2a in PAML. Nucleotide position refers to the position in the alignment following trimming – original coordinates of the chicken sequence can be found in Table 4B; Suppl. Data. (B) Using the same alignment, the free-ratios model in PAML was implemented to determine along which branches positive selection has been strongest. Each branch is labelled with its respective dN/dS value, and branches are coloured according to dN/dS.

To determine which lineage(s) have undergone positive selection in avian phylogeny, the same alignment was analysed using the ‘free-ratios’ test in PAML, which assigns a ω value for each branch in the tree. The most striking burst of positive selection was found at the base of the Galliformes, consistent with a change of ZAP function in this lineage (Fig. 5B). Weaker instances of positive selection were found in the Neoaves, but purifying selection predominated elsewhere in the tree. As adaptive evolution frequently associates with genes involved in host-viral arms races (39, 52), the findings are consistent with rapid host evolution of the ZAP gene in response to RNA virus-induced selection of ZAP-associated antiviral functions within the 150 million year period of avian diversification.

### Composition of mammalian and avian interferon-related genes

If ZAP differentially regulated expression of IFNs, ISGs and non-ISGs through differences in gene composition, then there should be differences in their dinucleotide compositions in avian species that possessed or lacked functional responses to high CpG replicons and viruses investigated in the study. Formally, CpG suppression should be more marked in duck transcriptomes compared to those of chicken based on the observed functional differences in ZAP (Figs. 2, 3). It is further possible that genes preferentially expressed during an antiviral response, such as interferon-stimulated genes and type 1 interferons, may be under more stringent selection against CpG and UpA as they may need to function in cells with elevated levels of ZAP expression.

To investigate these possibilities, sequence compositions of coding region sequences of mRNAs of *G. gallus* (non-restrictive) and *A. platyrhynchos* (restrictive) were compared with each and with those of human mRNAs using non-redundant RefSeq entries. mRNA sequences from each species showed consistent suppression of CpG and UpA representation, to extents related to their G+C composition (Figs. 6, 7). By linear regression, G+C content was associated with 38%-50% of the variability in CpG representation and 25%-28% of UpA (based on R^2^ values). Relationships, expressed as values of m and c in the linear regression relationship (freq. dinucleotide) = (m * G+C content) + c were highly similar between host species, although formal comparison of regression values through calculation of t values^1^ identified significant differences in trajectories between species (Table S6, Fig. S2; Suppl. Data). However, while there was functional restriction of high CpG E7 replicons and IAV in the duck-derived CCL-141 cell line but none was detectable in the DF-1 cell line from chickens, cellular mRNA showed a slightly higher CpG representation in duck cells, whereas the expectation would be for the duck transcriptome to show greater CpG suppression if its composition were conditioned by ZAP activity.

**FIGURE 6.**
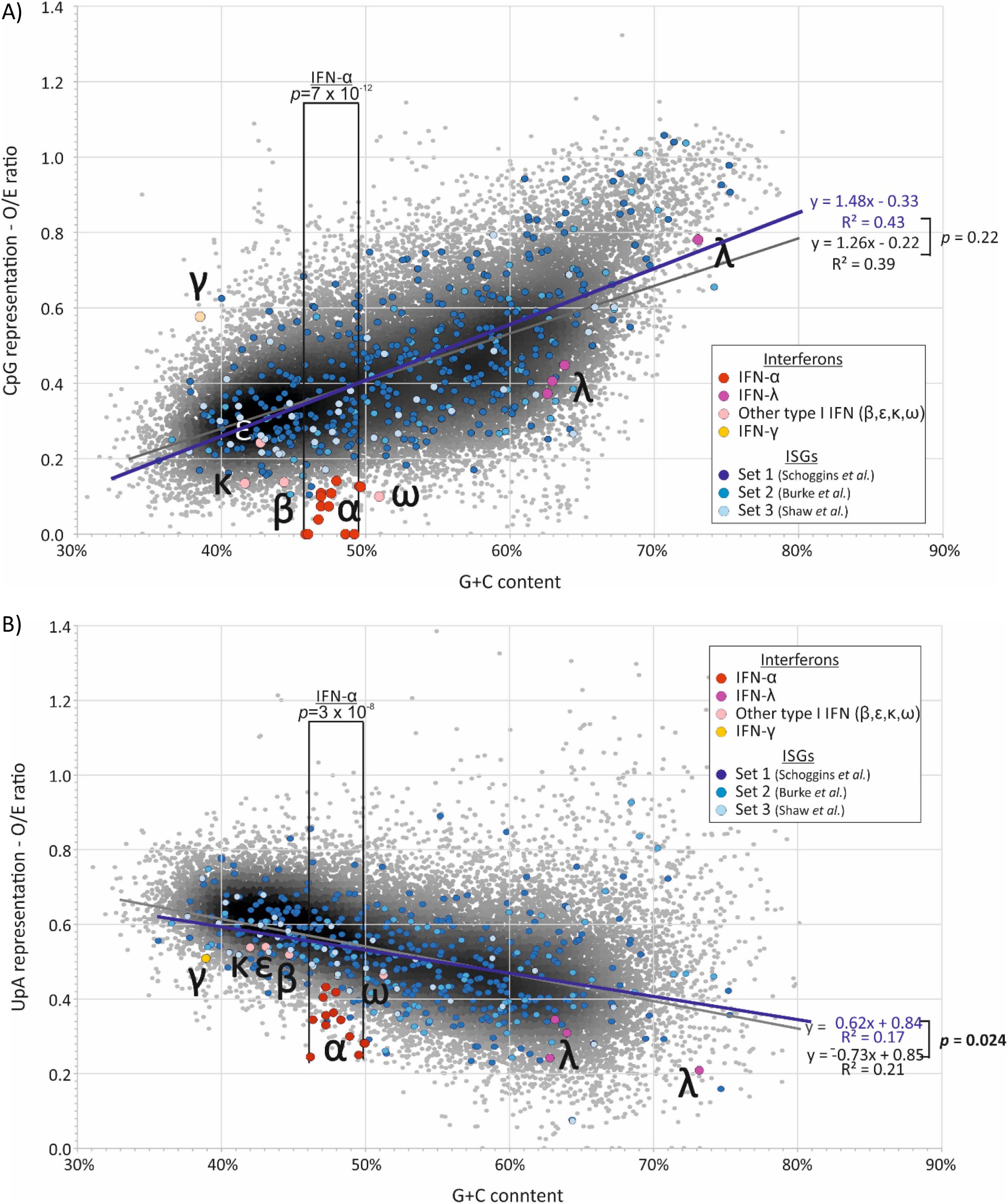
COMPARISON OF DINUCLEOTIDE FREQUENCIES OF HUMAN ISGs AND IFN GENES WITH THE BULK HUMAN mRNA TRANSCRIPTOME A comparison of (A) CpG and (B) UpA dinucleotide representation in human ISGs and interferon genes with the bulk human mRNA transcriptome (of coding sequence lengths ≥450 bases). CpG and UpA representations (observed frequency / frequency expected base on mononucleotide frequencies) of IFN-α paralogues were compared with human mRNA sequences in the G+C content range spanning the former (40.2%-50.0%) using an independent samples t-test. CpG and UpA representations in ISGs and mRNA were compared using regression analysis (see Results text; values for individual ISG datasets are shown in Table 2). Sources of ISG sequences: Set 1: (53); Set 2: (54); Set 3: (38).

**FIGURE 7.**
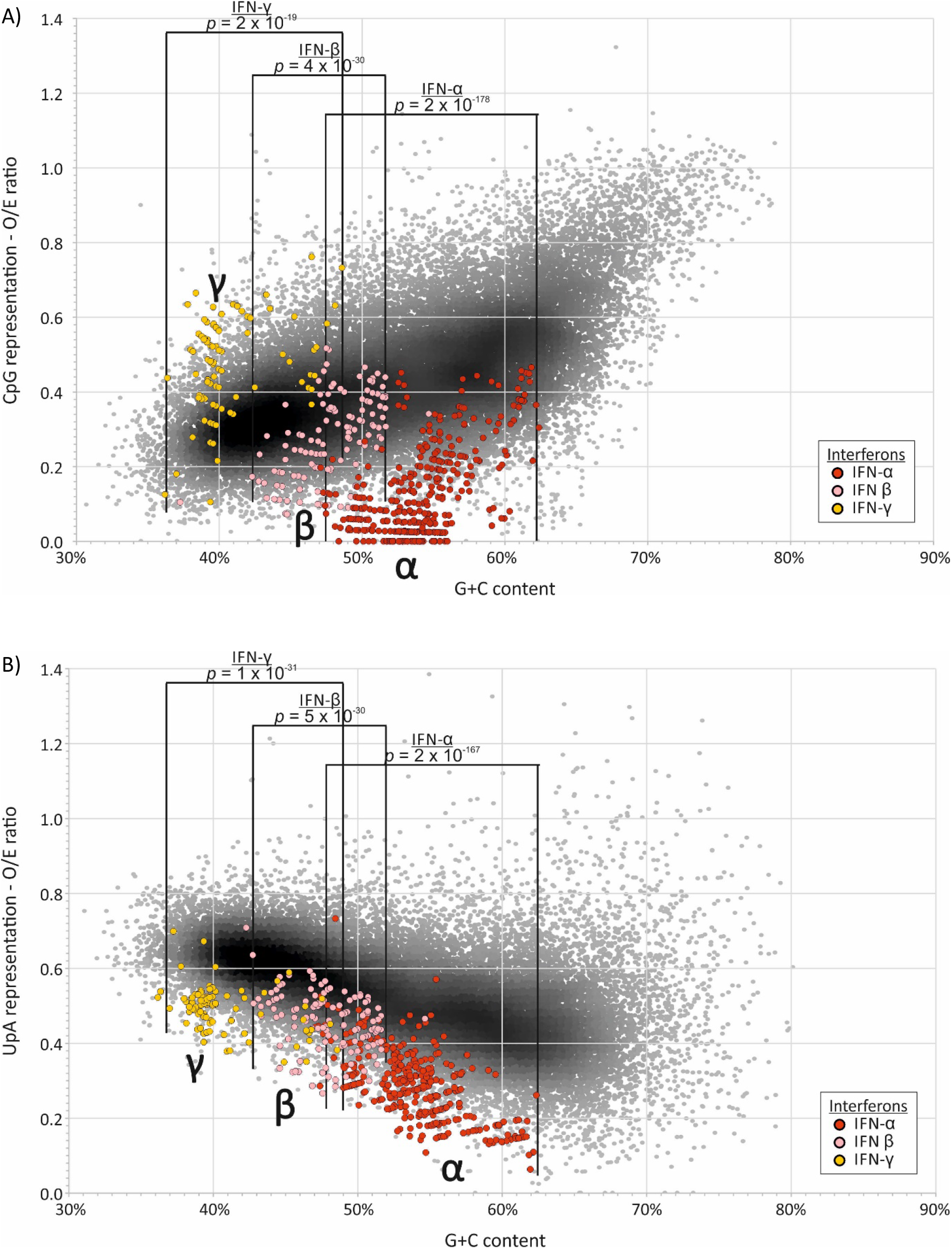
CpG AND UpA REPRESENTATION IN MAMMALIAN INTERFERON GENES Comparison of dinucleotide representations of IFN-α, IFN-β and IFN-γ in mammalian species with the bulk human mRNA transcriptome (IFN gene accession numbers and host species are listed in Table S8; Suppl. Data). Differences in values over defined G+C content ranges specific for each IFN gene class were analysed using the independent sample paired t test.

Similar comparisons were made between mRNA composition of each species with their corresponding subsets of genes identified as ISGs. For human genes, lists of upregulated genes on IFN stimulation were obtained from three separate published sources and analysed collectively (Figs. 6) or individually for composition or regression comparisons (Table 1; Table S7; Suppl. Data). As a group, human ISGs showed similar mean values in CpG or UpA representation and no significant differences from sequences in the wider human mRNA transcriptome (Table 1A). Regression analysis similarly revealed no significant difference in relationships between G+C composition and either CpG or UpA representation (Table 1B). On analysing individual subsets of ISGs, ISGs reported by Schoggins *et al.* (53) and ISGs upregulated by poly(I:C) treatment (54) showed a comparable range of G+C and dinucleotide representations to those of total mRNA sequences (Table 1A). However, the subset of 50 most upregulated ISGs reported by Shaw *et al.* (38) showed a mean CpG O/E ratio of 0.35, significantly lower than the mean of 0.45 of human mRNAs (*p* = 0.0003). However, the ISGs also showed a lower mean G+C content (49.7% compared to 52.9%; *p* = 0.017). Regressions of this ISG subset and human mRNAs were comparable (*p* = 0.45; Table 1B).

**TABLE 1.** COMPARISON OF BASE AND DINUCLEOTIDE COMPOSITIONS OF IFN-STIMULATED GENES (ISGs) AND IFN-REGULATED GENES (IRGs) WITH TOTAL

| Dataset | Total<sup>1</sup> | G+C<sup>2</sup> | *p*<sup>3</sup> | CpG<sup>2</sup> | *p*<sup>3</sup> | UpA<sup>2</sup> | *p*<sup>3</sup> |
| --- | --- | --- | --- | --- | --- | --- | --- |
| Human mRNAs | 35982 | 52.9% |  | 0.4416 |  | 0.5207 |  |
| Human ISGs | 421 | 52.9% | (*p*=0.93) | 0.4514 | (*p*=0.46) | 0.5142 | (*p*=0.20) |
| Burke *et al.*<sup>6</sup> | 101 | 54.1% | (*p*=0.20) | 0.4509 | (*p*=0.75) | 0.4959 | (*p*=0.029) |
| Schoggins *et al.*<sup>7</sup> | 401 | 52.6% | (*p*=0.54) | 0.451 | (*p*=0.41) | 0.5063 | (*p*=0.12) |
| Shaw *et al.*<sup>8</sup> | 47 | 49.7% | (*p*=0.017) | 0.3544 | (*p*=3×10<sup>-4</sup>) | 0.5103 | (*p*=0.72) |
| Human IRG | 47 | 55.21% | (*p*=0.12) | 0.5014 | (*p*=0.062) | 0.5253 | (*p*=0.78) |
| Chicken mRNAs | 49888 | 51.50% |  | 0.463 |  | 0.524 |  |
| Chicken ISGs | 215 | 51.70% | (*p* = 0.642) | 0.492 | (***p* = 0.013**) | 0.535 | (*p*=0.423) |
| Shaw *et al.*<sup>9</sup> | 41 | 49.80% | (*p* = 0.246) | 0.452 | (*p* = 0.955) | 0.53 | (*p*=0.883) |
| Chicken IRG<sup>9</sup> | 41 | 56.49% | (***p*=3.0E-5**) | 0.576 | (***p* = 5.0E-5**) | 0.493 | (*p*=0.064) |
| Duck mRNAs | 37925 | 50.60% |  | 0.472 |  | 0.549 |  |
| Duck ISGs | 212 | 50.10% | (*p*=0.58) | 0.492 | (*p*=0.069) | 0.558 | (*p*=0.059) |

B) Regression analysis
| Dataset | Total<sup>1</sup> | CpG RSq<sup>4</sup> | *p*<sup>5</sup> | UpA RSq<sup>4</sup> | *p*<sup>5</sup> |
| --- | --- | --- | --- | --- | --- |
| Human mRNAs | 35982 | 0.39 |  | 0.249 |  |
| Human ISGs | 421 | 0.429 | (*p*=0.22) | 0.166 | (*p*=0.023) |
| Burke *et al.*<sup>6</sup> | 101 | 0.52 | (*p*=0.72) | 0.06 | (*p*=0.03) |
| Schoggins *et al.*<sup>7</sup> | 401 | 0.43 | (*p*=0.25) | 0.23 | (*p*=0.49) |
| Shaw *et al.*<sup>8</sup> | 47 | 0.782 | (*p*=0.45) | 0.423 | (*p*=0.20) |
| Human IRGs<sup>9</sup> | 47 | 0.574 | (*p*=0.69) | 0.063 | (*p*=0.03) |
| Chicken mRNAs | 49888 | 0.48 |  | 0.25 |  |
| Chicken ISGs | 215 | 0.52 | (*p* = 0.99) | 0.22 | (***p*=0.001**) |
| Shaw *et al.*<sup>9</sup> | 41 | 0.31 | (*p* = 0.52) | 0.49 | (***p* = 0.04**) |
| Chicken IRG<sup>9</sup> | 41 | 0.6 | (*p* = 0.87) | 0.25 | (*p* = 0.92) |
| Duck mRNAs | 37925 | 0.5 |  |  |  |
| Duck ISGs | 212 | 0.6 | (*p*=0.51) | 0.28 | (*p*=0.49) |

<sup>1</sup>Number of coding region sequences >450 bases in length analysed
<sup>2</sup>Mean G+C content or dinucleotide O/E ratios of sequences
<sup>3</sup>Significance of differences in base or dinucleotide frequencies between selected ISG/IRG subsets and total transcriptome mRNA sequences, evaluated by Mann-Whitney U test. Cells shaded by probability
range
<sup>4</sup>Correlation coefficient between G+C content and CpG or UpA O/E ratios
<sup>5</sup>Comparison of regressions between total transcriptome and ISG or IRG subsets. Cells shaded by probability range
<sup>6, 7, 8</sup>Sources in refs. (54), (64) and (38) respectively
<sup>9</sup>Interferon-regulated genes

We obtained similar data for avian ISGs identified as homologues of mRNA sequences of ISGs from the three human ISG datasets (Table S8; Suppl. Data). G+C contents of chicken ISGs similarly spanned the range of host mRNA sequences (Fig. 8), with no significant differences in CpG or UpA representation in these or ISG homologues in ducks (Table 1A), apart from a slightly elevated frequency of UpA dinucleotides in duck-derived ISGs. In a separate analysis, the 50 most upregulated ISGs in chicken (listed in (38)) similarly showed no significant difference in CpG representation from chicken mRNAs, either by composition comparison or by regression. Both CpG and UpA representations were comparable with host transcriptome and with each other by on G+C composition regression analysis (Fig. 7, Table 1B; Table S6, Suppl. Data). Interferon-regulated genes (IRGs) showed significantly higher CpG and UpA compositions, but also a higher G+C content, and there was no significant difference in their distributions by linear regression comparisons.

**FIGURE 8.**
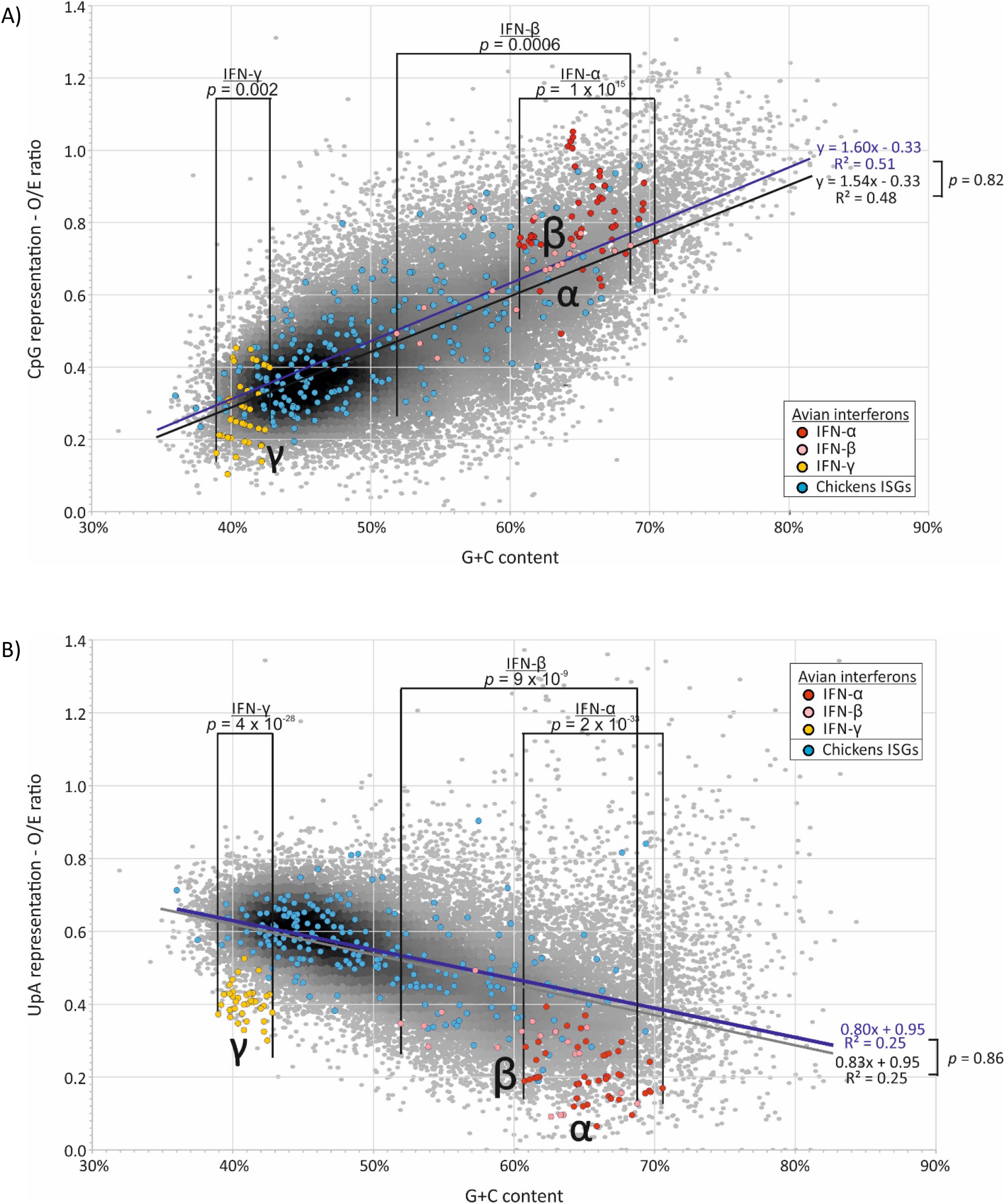
CpG AND UpA DINUCLEOTIDE REPRESENTATION IN THE CHICKEN TRANSCRIPTOME, ISGs AND AVIAN INTEFERON-α AND −γ GENES A comparison of (A) CpG and (B) UpA dinucleotide representation in chicken ISGs and avian interferon genes (listed in Table S8, S10; suppl. Data) with chicken mRNA transcriptome sequences (of coding sequence lengths ≥450 bases). Chicken ISGs were identified through homology searching for homologues of human ISGs listed in (38, 53, 54) (Table S8; Suppl. data). Differences in dinucleotide representations over defined G+C content ranges specific for each IFN gene class were analysed using the independent sample paired t test.

In contrast to cellular mRNAs and ISGs, IFN genes (listed in Table S9; Suppl. Data) showed often marked dinucleotide frequency differences from the host transcriptome. Formal analysis of these differences was complicated by the restricted ranges of G+C contents compared to that of the host transcriptome. Accordingly, frequency comparisons were performed within the G+C content range of the IFN subset being compared, for example between 40.2% and 50% for the comparison of human IFN-α genes with host sequences (Fig. 7; Table 2). Summarising, human and mammalian IFN-α showed highly suppressed frequencies of both CpG and UpA compared to cellular genes of similar G+C contents (Figs. 6, 7; Table 2). IFN-β genes showed lower degrees of suppression of CpG, while IFN-γ gene CpG frequencies were higher than those of the corresponding mRNAs. Both IFN-β and IFN-γ genes showed marked suppression of UpA frequencies.

**TABLE 2.** MEAN CpG AND UpA FREQUENCIES IN HUMAN, MAMMALIAN AND AVIAN INTERFERON GENES

| Host | Gene | Mean CpG O/E |  |  | Mean UpA O/E |  |  |
| --- | --- | --- | --- | --- | --- | --- | --- |
|  |  | mRNA<sup>1</sup> | Subset | *p*<sup>2</sup> | mRNA<sup>1</sup> | Subset | *p*<sup>2</sup> |
| Human | IFN-α | 0.352 | 0.064 | $6 \times 10^{-12}$ | 0.588 | 0.347 | $3 \times 10^{-8}$ |
| Mammalian | IFN-α | 0.449 | 0.140 | $2 \times 10^{-178}$ | 0.491 | 0.275 | $2 \times 10^{-167}$ |
| | IFN-β | 0.396 | 0.298 | $4 \times 10^{-20}$ | 0.542 | 0.447 | $5 \times 10^{-30}$ |
| | IFN-γ | 0.330 | 0.478 | $9 \times 10^{-19}$ | 0.607 | 0.486 | $1 \times 10^{-30}$ |
| Avian | IFN-α | 0.665 | 0.809 | $1 \times 10^{-15}$ | 0.408 | 0.203 | $2 \times 10^{-33}$ |
| | IFN-β | 0.562 | 0.666 | 0.0006 | 0.444 | 0.269 | $9 \times 10^{-9}$ |
| | IFN-γ | 0.325 | 0.277 | 0.002 | 0.616 | 0.406 | $4 \times 10^{-28}$ |

<sup>1</sup>Mean CpG or UpA O/E ratios for mRNA sequences within the G+C content range of the ISG subset.<sup>2</sup> Significance of differences in dinucleotide frequencies between interferon genes and total transcriptome mRNA sequences, evaluated by Mann-Whitney U test. Cells shaded by probability range

In contrast to mammalian interferons, avian IFN-α and IFN β genes (listed in Table S10; Suppl. Data) showed elevated CpG representations compared to chicken (Fig. 8, Table 2) and duck (data not shown) mRNA transcriptomes, although there was a moderate suppression of CpG dinucleotides in IFN-γ genes. In marked contrast, all type I and II IFN genes showed significantly suppressed frequencies of UpA compared to the avian cellular transcriptome (Fig. 8; Table 2).

### CpG and UpA representation in avian and mammalian RNA viruses

RNA viruses with +strand and - strand genomes infecting vertebrates and retroviruses typically display substantial suppression of CpG and UpA dinucleotide frequencies (25–27). Representation of both dinucleotides and their G+C dependence are comparable to those observed in cellular mRNA sequences (25). As ZAP is considered to possess a primarily antiviral function, suppression of CpG and UpA dinucleotides of viruses infecting different hosts may be influenced by potential differences in ZAP activity in different avian and mammalian cells.

To investigate this, we collected datasets of RNA viruses from the ICTV virus metadata resource (VMR) representative of each species of + and – strand RNA viruses and retroviruses annotated for mammalian or avian hosts but excluding dual-host viruses such as arboviruses (Table S11; Suppl. Data). Genomes were split into component genes and compositional analyses performed on their coding sequences; sequences shorter than 450 bases in length were excluded from analysis. Overall, a total of 1929 genes from 554 mammalian RNA virus genomes and 319 genes from 74 avian viruses drawn from 21 different virus families were compared (Tables S11, S12; Suppl. Data). In general, distributions of CpG and UpA frequencies in RNA viruses infecting both mammals and birds were fully overlapping with each other and with dinucleotide compositions of cellular genes (Fig. 9). Although the mean G+C content of RNA viruses was lower than that of human or avian mRNA sequences, suppression of CpG and UpA dinucleotides showed a similar relationship with G+C content as observed in cellular mRNAs. Correlation coefficients and trajectories of G+C / dinucleotide relationships for mammalian and avian RNA viruses were minimally different from each other and comparable to those of avian (Fig. 9B) and mammalian (Fig. 9A) mRNA genes.

**FIGURE 9.**
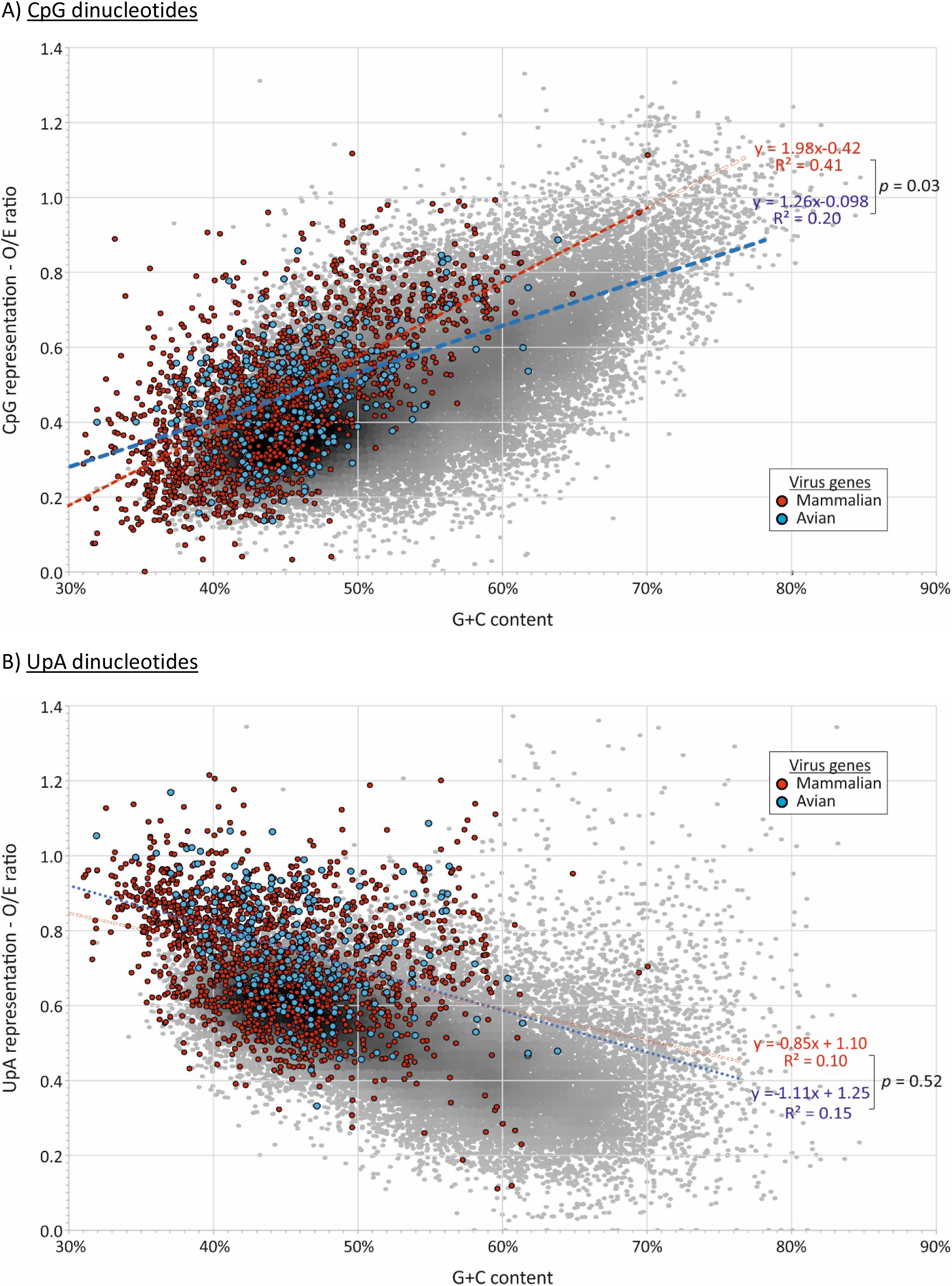
COMPARISON OF CpG AND UpA COMPOSITION IN RNA VIRUSES INFECTING MAMMALS AND BIRDS Observed / expected ratios of (A) CpG and (B) UpA RNA viruses (listed in Table S11; S12, Suppl. data) infecting mammals and birds. Ratios have been overlaid on values for avian (chicken) mRNA sequences to provide context. Analysis restricted to coding sequences of length >450 bases.

Because viruses assigned to different virus families may show systematic differences in composition (particularly G+C content), another way to compare effects of host is to compare compositions of avian and mammalian viruses assigned to the same family (Fig. 10A; Table S13A; Suppl. Data). Differences in G+C content, CpG and UpA representations were observed between the 10 virus families with mammalian and avian members (excluding orthomyxoviruses). However, there was no consistency in whether CpG or UpA were over- or under-represented in mammalian compared to avian viruses between them; combining the data from all RNA viruses, net differences in all three composition metrics were close to zero.

**FIGURE 10.**
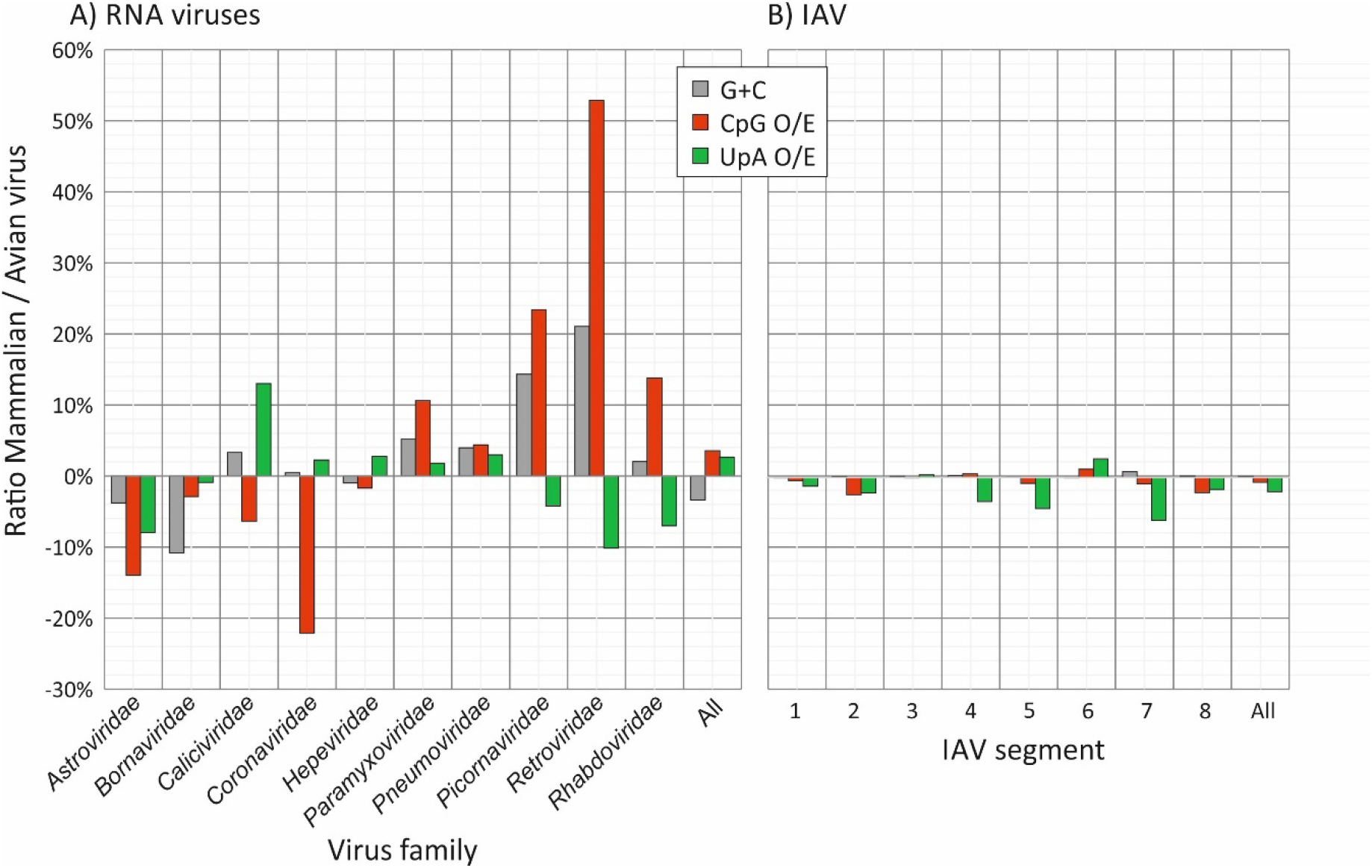
DIFFERENCES IN G+C CONTENT AND CpG AND UpA REPRESENTATION IN GENE SEQUENCES OF RNA VIRUSES INFECTING MAMMALS AND BIRDS Comparison of virus composition between viruses infecting mammals and birds, expressed as fractional differences in G+C content and CpG and UpA dinucleotide representation calculated as:

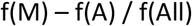

where f(M) is the mean composition of mammalian viruses, f(A) of avian viruses, and f(All) is the overall mean. Analyses were performed separately for different virus families and for different segments of flu strains isolated from mammalian (human) and avian sources (duck, chicken). Further analyses of differences between duck and chicken hosts, analyses where recently zoonotic H5N1 strains have been excluded and CpG and UpA representations calculated independently of protein coding are provide in Tables S15 (Suppl. Data). Source sequences are listed in Tables S10 and S11 (Suppl. Data).

Influenza A virus (IAV) is an example of a multi-vertebrate host virus, and serotypes infecting a range of avian and mammalian host species have been extensively characterised genetically. Ducks are regarded as the primary reservoir of IAV worldwide, although the virus will transmit readily to other avian species and to mammals. Adaptation to a new host, such as the H1N1 serotype in human following its pandemic spread from 1918 (55) has been proposed to drive reductions of CpG dinucleotide frequencies (35, 56) potentially associated with a more active ZAP-mediated recognition of avian-adapted viruses (40). To investigate whether differences in ZAP function in ducks, chickens and mammalian cells led to systematic differences of CpG and UpA representation, we analysed dinucleotide compositions in a large dataset of representative isolates of IAV with annotated hosts from the Influenza Research Database (mammalian: 2949 isolates, primarily H1N1 and H3N2 serotypes; avian: 6417 isolates; Table S14, S15; Suppl. Data). Coding region sequences in each segment were analysed separately (Fig. 10B).

As observed in the analysis of different virus families, there were (smaller) differences in CpG and UpA composition of IV strains infecting humans and birds between different segments although there was no consistent directional trend. Combining data for the whole genome, there was a net difference of +1% difference in G+C content, +1.0% in CpG and −1.5% in UpA representation in IAV strains infecting mammals compared to birds (Fig. 10B; Table S13C; Suppl. Data). Similar results were obtained on repeating the analysis after exclusion of H5N1 strains (Table S13D; Suppl. Data); these have recently zoonotically spread into humans and which may therefore preserve genome compositions more typical of avian variants. Finally, the analysis was repeated using CpG and UpA expected / observed ratios that considered only bases that could vary synonymously (corrected frequencies in SSE v.1.4; (44)) and which therefore removed potential biases originating from amino acid compositions of viral genes and the presence of fixed dinucleotides within specific codons. This again demonstrated no consistent host-associated differences between IAV segments in their CpG or UpA representations (Table S15A; Suppl. Data).

Since ducks and chicken cells varied in their restriction of CpG-high IAV variants or E7 replicons, we performed a further set of comparisons between IAV strains isolated from these two host species (Table 13B; Suppl. Data). No consistent differences in G+C contents was observed, although most segments showed marginally lower CpG and UpA representation in duck derived IAV isolates compared to those of chicken (mean for all segments: −0.64% and −2.0% for CpG and UpA respectively). Similar findings were obtained using coding corrected dinucleotide representations (Table S15B; Suppl. Data).

Overall, this analysis did not show convincing differences in dinucleotide compositions between mammalian and avian RNA viruses, nor between those infecting avian hosts with demonstrated differences in ZAP activity.

## DISCUSSION

This study investigated the extent to which physiological differences in ZAP activity observed between cell lines derived from mammals and bird may be reflected in their genome compositions. Of particular interest is the possible effect of ZAP restriction on genes expressed at high level during an antiviral response when ZAP was more active. Observed differences in ZAP-mediated restriction against CpG- and UpA-enriched replicons and viruses between cell lines from these different hosts provided the opportunity to investigate the relationship between ZAP activity and potential differences in depletion of CpG and UpA dinucleotides in cytoplasmic RNA sequences and in the viruses that infected them. While all mammalian cell lines investigated potently restricted CpG-high mutants of E7, avian cells showed variable inhibition of CpG-high E7 and IAV mutants, most apparent in duck cell lines (order Anseriformes) but not in cells from chicken or other galliforms. Avian cells further differed to mammals by their inability to restrict replication of mutants with elevated UpA frequencies. If ZAP shaped genome composition, then we might expect to observe consistent differences in CpG and UpA representation in host cell transcriptomes, particularly in ISGs and IFN genes where ZAP is upregulated during an antiviral response. Our findings of compositional equivalence in each of these gene sets in hosts with radically different ZAP activities is reviewed in the light of previously published data on compositional differences in a wide range of genes and host species.

### Host-associated differences in ZAP function

We documented substantial variability in the abilities of cell lines derived from different avian species to preferentially restrict the replication of high CpG mutants of IAV and the E7 replicon. Duck derived cells restricted high CpG IAV and E7 equally potently as the range of mammalian cell lines tested (Fig. 2), the pigeon cell line showed a marginal restriction of the high CpG E7 replicon, while all cell lines derived from chicken, other galliforms and zebra finch displayed none. It should be noted, however, that the varying permissivity of the different cell lines for E7 replication may originate from any number of cellular mechanisms and this required us to normalise replication to that of the WT replicon (y-axis; Fig. 2).

This format contrasts with a previous investigation of the activity of avian-derived ZAPs, where the replication of CpG modified mutants of HIV-1 was compared in mammalian HEK293T ZAP k/o cells co-transfected human/test species ZAP chimaeras and a cognate TRIM25 co-factor required for activity (40). Using this human cell transfection system, duck and eagle ZAP/TRIM25-transfected cells mediated marked and selective suppression of CpG-high mutants of HIV-1, consistent with the restriction in CCL-141 (duck) cells observed in the current study. However, while chicken ZAP/TRIM25-transfected cells also showed some antiviral activity, restriction was non-selective and similarly active in both CpG-H and WT variants of HIV-1. As our assay readout assigned restriction by normalisation to the replication of WT replicon, such non-selective activity in for example, the DF-1 chicken cell line, would have been undetectable. However, if we equate CpG-high/WT replication ratios (Fig. 2) with ZAP/TRIM25 selectivity (40), the avian species-associated differences we found were comparable to those identified in the heterologous expression system; specifically, chicken, turkey and zebra finch showed little CpG selectivity in virus inhibition, in contrast to duck and eagle-derived ZAP/TRIM25.

Additional evidence for host differences in dinucleotide-mediated virus recognition was provided by the absence of restriction of UpA-high E7 replicons (Fig. 2B) and compositionally modified IAV (Fig. 3B) in the duck cell line CCL-141 that was otherwise capable of potent restriction of CpG-high mutants. This contrasts with substantial restriction of the UpA-H double mutant of E7, comparable to that of the CpG-H mutant observed in all mammalian cells (Fig. 2A). As the recognition mechanism of UpA-high mutants remains undetermined, and may indeed be mediated through a non-ZAP PRR (12), these findings do not necessarily implicate an altered substrate specificity of avian ZAPs. Measurement of the restriction of UpA-high mutants of E7 using the avian ZAP/TRIM25 transfection method would be informative in this respect.

More broadly, the finding of active although variable ZAP-mediated restriction in different bird species is consistent with the evidence for positive selection in the ZAP gene in the evolution of birds, comparable to that inferred in primate and wider mammalian evolution (39, 57). The evidence for intense selection at the base of the galliform lineage (Fig. 5) is particularly interesting and may help explain some of the phenotypes seen with cells derived from this group of birds. It is noteworthy that these ZAP-based differences add to other avian lineage-based differences that may explain patterns of viral susceptibility. For example, the loss of RIG-I in galliforms has been attributed to the differences in susceptibility of ducks and chickens to AIV as have the expression and response patterns of other PRRs (58, 59). Clearly, ZAP-mediated viral restriction pathways may be an important additional factor defining the outcome of viral infections in different species.

Irrespective of the underlying drivers of variability in ZAP function, these observations provide an opportunity to investigate whether ZAP plays a role in shaping the composition of cellular transcriptomes or of the RNA viruses that infect different hosts. Our analysis of cellular mRNA transcriptomes and the subset of interferon-stimulated genes that may have evolved to function in cells with high levels of ZAP expression, failed to find differences in CpG or UpA suppression between chicken and duck genomes. Collectively, avian genes showed comparable degrees of suppression of both dinucleotides to those observed in mammalian genomes. Our extended analysis of RNA virus genome composition further demonstrated no substantial differences between RNA viruses infecting mammals from those infecting birds, findings that contrast with previous conclusions (40, 60). Of direct relevance to the transmission of IAV between bird species, we found no evidence for systematic compositional differences between IAV strains isolated from ducks and chickens, even though we might hypothesise that the absence of ZAP-mediated discrimination of CpG-high sequences (40) in chicken cells might have enabled variants with less CpG-suppressed genomes to have evolved.

More striking is the more general similarity in CpG and UpA representations between mammalian and avian flu strains. It was previously shown following the zoonotic transmission of H1N1 from an inferred avian host into humans was associated with a sustained and decades long reduction in CpG dinucleotide frequencies as the H1N1 strain spread and became established in humans following its emergence in 1918 in the Spanish flu pandemic. (55). This was accounted for by the mechanistically unproven supposition that human cells were more restrictive for CpG than avian cells (56), that in retrospect might be interpreted as indicative of more active ZAP function (7, 40). It was further proposed that recently transmitted IAV strains may be more pathogenic in humans because of their greater CpG frequencies and consequent activation of responses that lead to immunopathology (35, 56). While superficially attractive as an explanation for the pathogenicity of recently transmitted flu strains, such as those with H5 and H7 HA segments (61), the whole framework might be challenged on several grounds. Firstly, on an extended comparison, we found no systematic compositional difference between IAV strains recovered from human and either species of bird using linear regression that accommodates the effects of G+C content on CpG composition (Fig. 11; Table S13, Suppl. data). Most human strains included in the comparison were the H1N1 and H3N2 serotypes, long established in human populations, while the duck and chicken strains were primarily avian serotypes also with likely long term host associations. The compositional similarities therefore did not arise simply from possible recent cross-species transmission as might be the case for H5N1. The second grounds for questioning the hypothesis is the previously discussed demonstration of an equally potent ZAP mediated restriction of high CpG viruses in duck cells (Fig. 2B) and evidence for similar restriction of CpG mutants of HIV-1 by the duck-derived ZAP construct (40). As ducks form the principal reservoir of IAV in nature (62), it is somewhat problematic to base a hypothesis on an assumption that IAV is less compositionally restricted in avian hosts than in humans.

**FIGURE 11.**
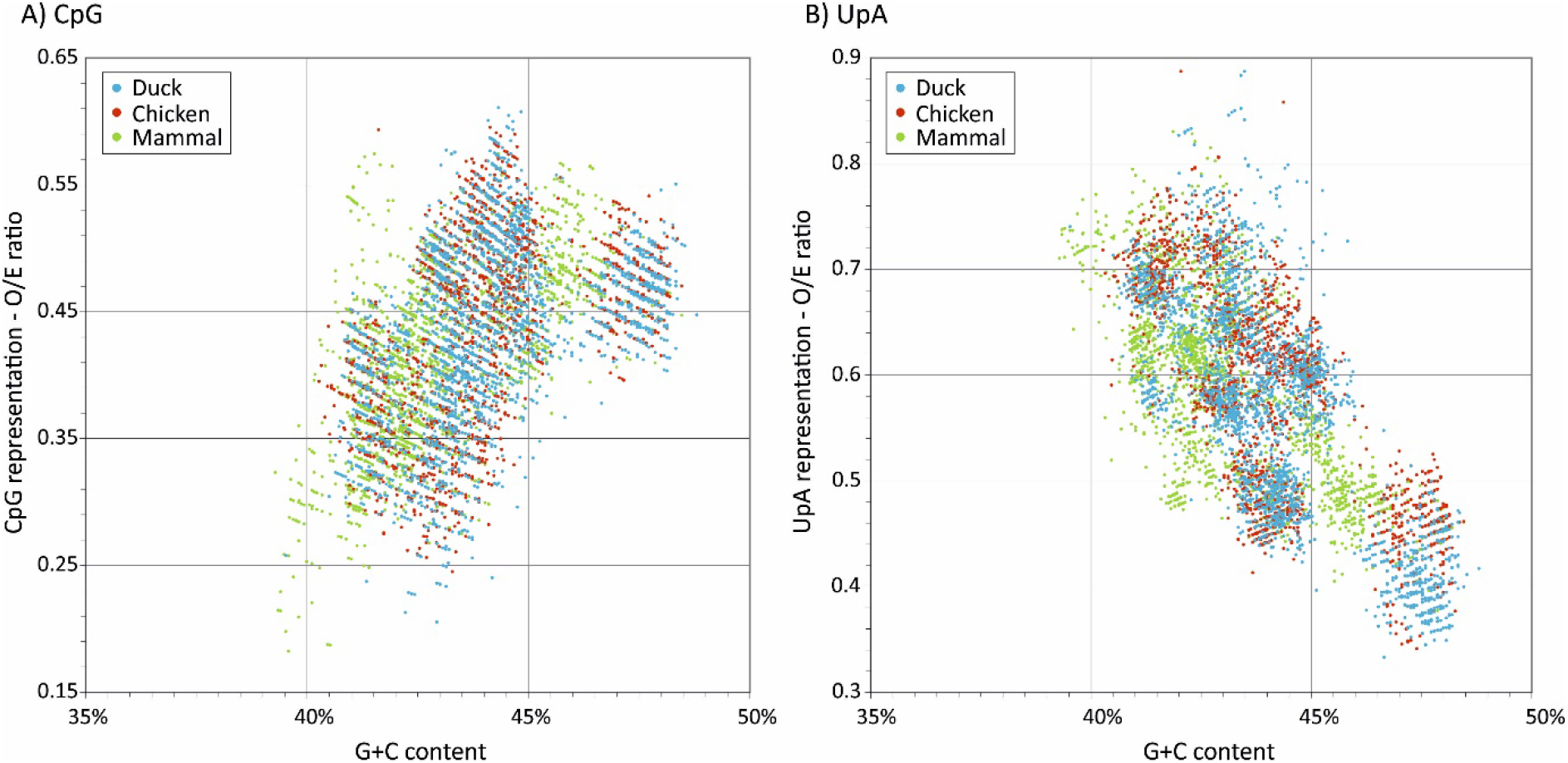
COMPARISON OF CpG AND UpA COMPOSITIONS OF MAMMALIAN AND AVIAN IAV STRAINS Distributions of CpG and UpA representations plotted against G+C content for IAV strains derived from different hosts; source sequences and serotype totals listed in Tables S13, S14 (Suppl. Data).

### Influence of ZAP on host transcriptome composition

The ability of ZAP to recognise and restrict RNA viruses appears to rely on its ability to detect clustered CpG dinucleotides (7). ZAP-mediated selection may be the mechanism underlying the previously inferred compositional selection against CpG and UpA suppression and their G+C compositional dependence arising from previous mutational modelling in mammalian transcriptomes (25). Further indirect evidence for the potential shaping effect of ZAP on host gene composition is provided by analysis of interferon genes. As previously described for human IFN-α genes (35, 63), human and all other mammalian type I IFN genes investigated showed substantially greater CpG suppression than other mRNA sequences (Fig. 7), but a similar dependence on G+C content as found in other mRNA sequences. We also observed much greater suppression of UpA in IFN genes compared together cellular mRNAs.

Long before the discovery of ZAP-mediated recognition of CpG dinucleotides, Greenbaum *et al.* proposed that this extreme suppression may enable greater of expression of IFNs perhaps at the expense of other cellular genes not engaged in antiviral defence (35, 56). One might develop the idea further in considering the possible effects of high-level expression of ZAP in interferon-induced cells on virus infection. ZAP over-expression may provide the mechanism for differential gene expression from genes specifically depleted for CpG and UpA dinucleotides. This form of gene regulation may contribute to the antiviral state and upregulated by virus infection. Hence, ZAP may enable cellular translation to be diverted towards interferon production as the rest of the transcriptome is regulated by increased degradation the virus-infected cells.

Extending this concept to other genes involved in the antiviral response is problematic. It was reported that CpG frequencies were lower in the subset induced by immune stimulation of mouse dendritic cells (35). Similarly, Shaw *et al.* have recently demonstrated significantly lower CpG compositions in the 50 most IFN-upregulated genes in human cells that unregulated or IFN downregulated genes (38) leading to their conclusion that ZAP may function as a cellular regulator of gene expression, enabling preferential expression of antiviral effector genes during the antiviral state when ZAP expression is upregulated and functionally more active. However, neither analysis acknowledged the effect of G+C content; re-analysis of the Shaw dataset in the current study showed a systematically lower G+C content than the average for cellular mRNAs and indeed frequencies of CpGs were not systematically different from expected values once this is taken into account by regression analysis (Table 1B). For each subset of human ISGs and their homologues in birds analysed in the current study, the identified upregulated genes fall in a similar trajectory to cellular mRNAs, and the comparable CpG and UpA compositions of ISGs from humans (and homologous genes in avian genomes) within the relevant G+C content ranges.

Overall, our extended analyses reach the rather negative conclusion that the activity of ZAP and other potential PRR recognising RNA sequences with abnormal dinucleotide compositions do not measurably lead to differences in cellular transcriptome compositions of human and avian cells. While there is profound suppression of CpG and UpA frequencies in mammalian type 1 IFN genes, this does not fully extend to those of avian homologues even though ZAP was active and discriminatory against CpG-enriched RNA sequences in ducks in this and previous studies (40). Furthermore, while neither UpA-enriched E7 or IAV mutants were restricted in duck or any other avian cell type, UpA frequencies were globally suppressed in both type I and type II (IFN-γ) IFN genes. There was no systematic difference in CpG and UpA frequency representation between mammalian and avian RNA virus genes, nor between IAV strains recovered from ducks and chickens, despite the functional differences in CpG and UpA restrictions in these different hosts. Overall, the study provides no evidence for a role of ZAP in differential regulation of gene expression, that might favour ISGs over other cellular genes in IFN-activated cells.

## ACKNOWLEDGEMENTS

We would like to thank Isabelle Dietrich from Pirbright institute, Ian Goodfellow, University of Cambridge, Paul Skinner from Animal Health Protection Agency and Rick Randall for generously providing cell lines. The work was funded by a Wellcome Investigator Award to PS (WT103767MA), Biotechnology and Biological Sciences Research Council (BBSRC) (Grant No. BB/M011224/1) for S.R.F. ALS has received support from the Department for Environment, Food and Rural Affairs (Grant No. OD0221) BBSRC (Grant No. BB/N023803/1 and BB/K004468/1) and the John Fell Fund (Oxford). ALS is a Jenner Investigator and has also received support from the Jenner Institute, Oxford.

## Footnotes

1 https://stats.idre.ucla.edu/spss/faq/how-can-i-compare-regression-coefficients-between-two-groups/

